# BioNetGMMFit: Estimating Parameters of a BioNetGen Model from Time-Stamped Snapshots of Single Cells

**DOI:** 10.1101/2022.12.08.519526

**Authors:** John Wu, William CL Stewart, Ciriyam Jayaprakash, Jayajit Das

## Abstract

**Background:** Mechanistic models are commonly employed to describe signaling and gene regulatory kinetics in single cells and cell populations. Recent advances in single-cell technologies have produced multidimensional datasets where snapshots of copy numbers (or abundances) of a large number of proteins and mRNA are measured across time in single cells. The availability of such datasets presents an attractive scenario where mechanistic models are validated against experiments, and estimated model parameters enable quantitative predictions of signaling or gene regulatory kinetics. To empower the systems biology community to easily estimate parameters accurately from multidimensional single-cell data, we have merged a widely used rule-based modeling software package BioNetGen, which provides a user-friendly way to code for mechanistic models describing biochemical reactions, and the recently introduced CyGMM, that uses cell-to-cell differences to improve parameter estimation for such networks, into a single software package: BioNetGMMFit.

**Results:** BioNetGMMFit provides parameter estimates of the model, supplied by the user in the BioNetGen markup language (BNGL), which yield the best fit for the observed single-cell, timestamped data of cellular components. Furthermore, for more precise estimates, our software generates confidence intervals around each model parameter. BioNetGMMFit is capable of fitting datasets of increasing cell population sizes for any mechanistic model specified in the BioNetGen markup language.

**Conclusion:** By streamlining the process of developing mechanistic models for large single-cell datasets, BioNetGMMFit provides an easily-accessible modeling framework designed for scale and the broader biochemical signaling community.

## 1 Introduction

Recent advancements in single-cell technologies have allowed for the measurement of cell-to-cell differences in mRNA/protein abundances [1–3]. These differences enable the evaluation of means and higher-order moments (e.g. variances and covariances), which can be utilized to improve the estimation of model parameters. One potential strategy, which is especially useful in our case where the probability distribution for the data (i.e. the *likelihood*) is unknown, is to estimate model parameters by minimizing the differences between sample moments and their corresponding predicted moments computed from *in silico* models. However, it is not obvious how any set of moment differences should be summarized, especially since sample moments (and hence moment differences) can vary considerably in single-cell data. The Generalized Method of Moments (GMM), widely used in econometrics [4, 5] and described in greater detail below, provides a systematic approach to incorporate means and higher-order moments in parameter estimation [6, 7]. Within GMM, the moment differences are combined into a single measure of cost (i.e. *distance*) using a system of weights that efficiently accounts for fluctuations across sample moments (see A.1 GMM Primer for more details). In practice, to find parameter values that minimize the GMM cost, one usually needs an optimization algorithm such as gradient-descent [8] or stochastic algorithms such as simulated annealing [9] or parallel tempering [10, 11]. In recent years, a class of meta-heuristic optimization algorithms such as Particle Swarm Optimization (PSO) that do not require the calculation of gradients and can be easily parallelized has been developed (see [12, 13] for a pedagogical review).

Rule-based modeling approaches, such as BioNetGen [14] and libRoadRunner [15], have been developed to address combinatorial complexity in modeling biochemical reactions in signaling and gene regulatory networks [16, 17]. These approaches provide a user-friendly way to construct models. Estimating model parameters, such as reaction rates, is crucial for improving model predictions and quantifying underlying mechanisms described by the model using experimental measurements. With the emergence of software packages such as PyBioNetFit [18], parameter estimation from rule-based models using bulk measurements of selected proteins (e.g. average or total protein abundances observed over time) has become possible.

We introduce BioNetGMMFit, a software tool that uses GMM to improve parameter estimation in BioNetGen models by exploiting the additional information in single-cell snapshot data. This tool requires users to supply a BioNetGen model .bngl file, time-stamped snapshot abundance data files, and run configuration files. BioNetGMMFit utilizes the GMM analysis of time-stamped protein abundances, as implemented in CyGMM [7], to estimate the parameters of the BioNetGen model, provide confidence intervals, predict moments at future times, and report the minimum cost (i.e., the distance between the sample moments and the moments predicted by the model using the GMM estimate of the parameters). While GMM can be used in conjunction with a wide variety of optimization routines, we use PSO to optimize our cost function because it does not require gradient calculations, scales well with higher-dimensional search spaces, has a relatively short run-time, and is easily parallelized on a compute cluster. Additionally, users can tune the PSO hyperparameters, which can affect the optimization’s efficiency and accuracy. As such, in addition to being written in C++, BioNetGMMFit is scalable with increasing data sizes on high-performance computing clusters. BioNetGMMFit is also available through Docker as C++ compilation varies across different operating systems. However, for improved performance, BioNetGMMFit can be compiled as a C++ executable through cmake and has been tested on Linux operating systems.

The manuscript is organized as follows. In the Software Description section, we provide detailed information on the implementation and use of BioNetGMMFit. Next, in the Results section, we present three examples of modeling single-cell data where we apply BioNetGMMFit and compare it with existing software tools. Finally, we offer our conclusions, discuss future directions and limitations in the Discussion section.

## 2 Software Description

To illustrate in detail how one might use BioNetGMMFit, a parameter estimation task with an experimental CD8+ T cell dataset was performed. As shown in Figure 1, BioNetGMMFit reads input data from several sources, including (i_1_) a directory containing a CSV file with initial protein abundances (X) and a directory containing CSV files with observed protein abundances at different time snapshots (Y); (i_2_) a time steps CSV file that contains the times at which the data of interest are observed (note that the data of interest may be a subset of the full data); (i_3_) a .bngl (BioNetGen) file that describes the mechanistic model capable of executing deterministic or stochastic kinetics; and (i_4_) a BioNetGMMFit hyperparameter configuration file that enables various features, such as generating time-series snapshot data from the BioNetGen model or changing the number of particles and steps of the PSO routine. A general overview of the BioNetGMMFit workflow is illustrated in Figure 2.

**Fig. 1.**
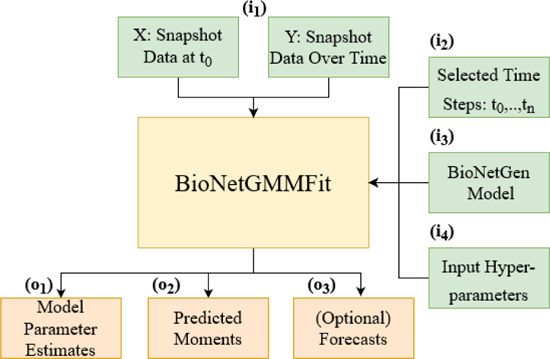
BioNetGMMFit Inputs and Outputs: The inputs and outputs to the program BioNetGMMFit (yellow box) are displayed. The green boxes, labeled with the letter “i”, are the inputs to the program. The two boxes included in i_1_ represent the input directories: X contains the initial conditions for the model and Y, all the observed snapshot data at different time points that must be passed to BioNetGMMFit. The box i_2_ is the time steps file that contains all the specified time points at which the concentrations are evaluated, by evolving the model given the initial conditions. i_3_ represents the model defined in BioNetGen containing the parameters to be estimated. The BioNetGen model (defined in BNGL) is used to evolve the initial conditions in X to fit the observed moments computed from the data in Y. i_4_ is the configuration file that contains the hyperparameters of the PSO (i.e the number of particles, steps, and PSO weights) that need to be defined by the user for the parameter estimation task. The orange boxes indicate outputs and are labeled by “o”. BioNetGMMFit produces two explicit outputs, o_1_, the parameter estimates of the BioNetGen model and o_2_, the corresponding time-evolved moments. The moments in o_2_ are generated using the parameter estimates that minimize the GMM cost between the observed moments and predicted moments. Lastly, the user can choose to forecast future trajectories of moments in o_3_.

**Fig. 2.**
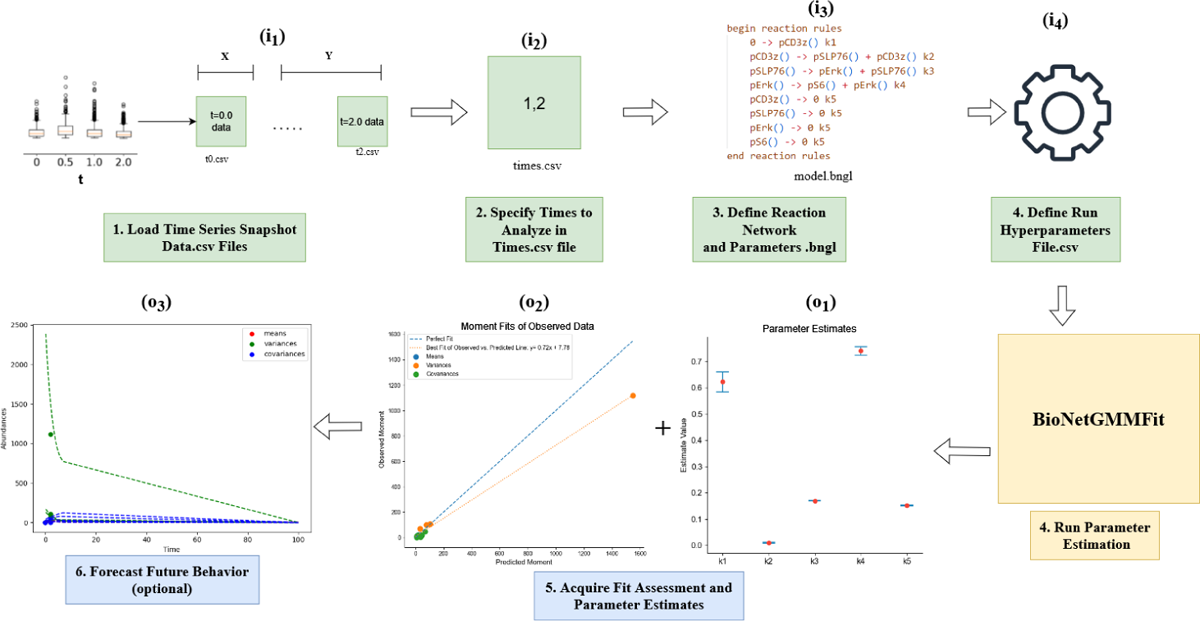
BioNetGMMFit Workflow: All “i”’s and “o”’s correspond to the same inputs and outputs defined in Figure 1. In this figure, we depict the steps needed to run BioNetGMMFit relating back to the inputs and outputs of Figure 1. To begin, the snapshot data of interest must be loaded by first organizing all .csv files into their corresponding initial conditions (X) and observed conditions (Y) directories, and then these (X) and (Y) directories (i_1_) must be specified to BioNetGMMFit. The black points in the data boxplot figure represents the outlier of the dataset (specifically CD8+ T cell dataset). Next, the user must define the time points that correspond to each of the snapshot files by creating and specifying a times.csv file (i_2_). Once all of the data is defined, the user must define and specify the BioNetGen model of interest (i_3_) to BioNetGMMFit. The last input required is the hyperparameters configuration file where the user must define all of the necessary parameters for a parameter estimation run, specifically related to the PSO algorithm that will be used to estimate the BioNetGen model parameters. Once all necessary inputs (i_1_, i_2_, i_3_, and i_4_) are defined, BioNetGMMFit will run its parameter estimation routine with GMM and PSO, returning parameter estimates (o_1_) (and confidence intervals if there are multiple replicate runs of PSO) as well as a *observed* = *predicted* line where observed moment values are on the y-axis and predicted moments (o_2_) generated from the estimated model parameter on the x-axis. Please note that the observed moments are calculated from the abundance data supplied by i_1_, and similarly the predicted moments are computed from the evolved abundances generated by simulating each initial sample with the BioNetGen model. Finally, while optional, users of BioNetGMMFit can predict moments of future time points (o_3_) of the BioNetGen model. Please refer to the Appendix for a more in-depth tutorial on how to use the command line version of BioNetGMMFit illustrated with the CD8+ T cell dataset.

When working with directories of data files, only one CSV data file is read from directory X to establish the initial set of protein abundances. Meanwhile, directory Y may contain multiple CSV files for each time step being analyzed. These data files are arranged such that each row contains the observations (i.e protein abundances) from an individual (or single) cell. To ensure that protein abundances or concentrations are non-negative, any single cell data containing negative protein values at a given time is removed before performing parameter estimation in BioNetGMMFit. Such a preprocessing step reduces the necessary steps the user must take to properly use BioNetGMMFit as negative values are commonly generated in CyTOF datasets during a processing step that spreads out zero readings in CyTOF measurements[19].

As an example, we will use a signaling kinetic model involving four proteins to analyze a CD8+ T cell CyTOF dataset. One biological question of interest that can be addressed with such data is to quantify the rate of the signal propagation between a pair of signaling proteins and study how the rates depend on the developmental state of the immune cell (e.g., naive vs memory CD8+ T cell). We applied BioNetGMMFit to describe signaling kinetics in CD8+ T cells stimulated by CD3 and CD28 antibodies where binding of the antibodies induce phosphorylation of the transmembrane CD3*ζ* chains which further lead to phosphorylation of the adaptor protein SLP76, MAPK kinase Erk and the ribosomal protein S6 following pCD3*ζ →* pSLP76 *→* pErk *→* pS6 [20]. We model the phosphorylation reactions shown above by first-order reactions. There are many intermediate biochemical reactions involved within the phosphorylation steps shown above and we assume the rates of the first-order reactions effectively capture the effect of those intermediate reactions. The phosphorylated protein species also go through degradation/ubiquitylation processes which are also approximated by first order decay reactions. In addition, pCD3*ζ* is assumed to be produced at a constant rate due to the signaling process. The model contains five rate constants, *θ*_1_*, · · ·, θ*_5_, which we estimate using BioNetGMMFit. BioNetGMMFit also plots confidence intervals as well as the observed and estimated moments as shown in Figure 3. The underlying datasets used in this parameter estimation task of the CD8+ T cells are from (https://dpeerlab.github.io/dpeerlab-website/dremi-data.html) [20]. From this dataset, one can compute time trajectories of the protein moments, means, variances, and covariances (see Figure 3). To make the presentation simple, we will only analyze trajectories at two time points: 1 minute and 2 minutes. In this case, 719 cells were measured at time point 1 minute and 653 cells were measured at time point 2 minute. Again, as shown in Figure 3, mechanistic model reaction rules are defined in the .bngl file where they are then converted into an SBML file using BioNetGen’s writeSBML() function as BioNetGen does not support its own C++ API. Using libRoadRunner’s C++ API [15], BioNetGMMFit converts this SBML file into a roadrunner object that is capable of simulating the defined model to user-specified time intervals. While there are many other reaction network simulators such as PySB [21], COPASI [22], and AMICI [23], libRoadRunner was primarily chosen for its speed and ease of customizability due to its function as a C++ API rather than a standalone software package.

**Fig. 3.**
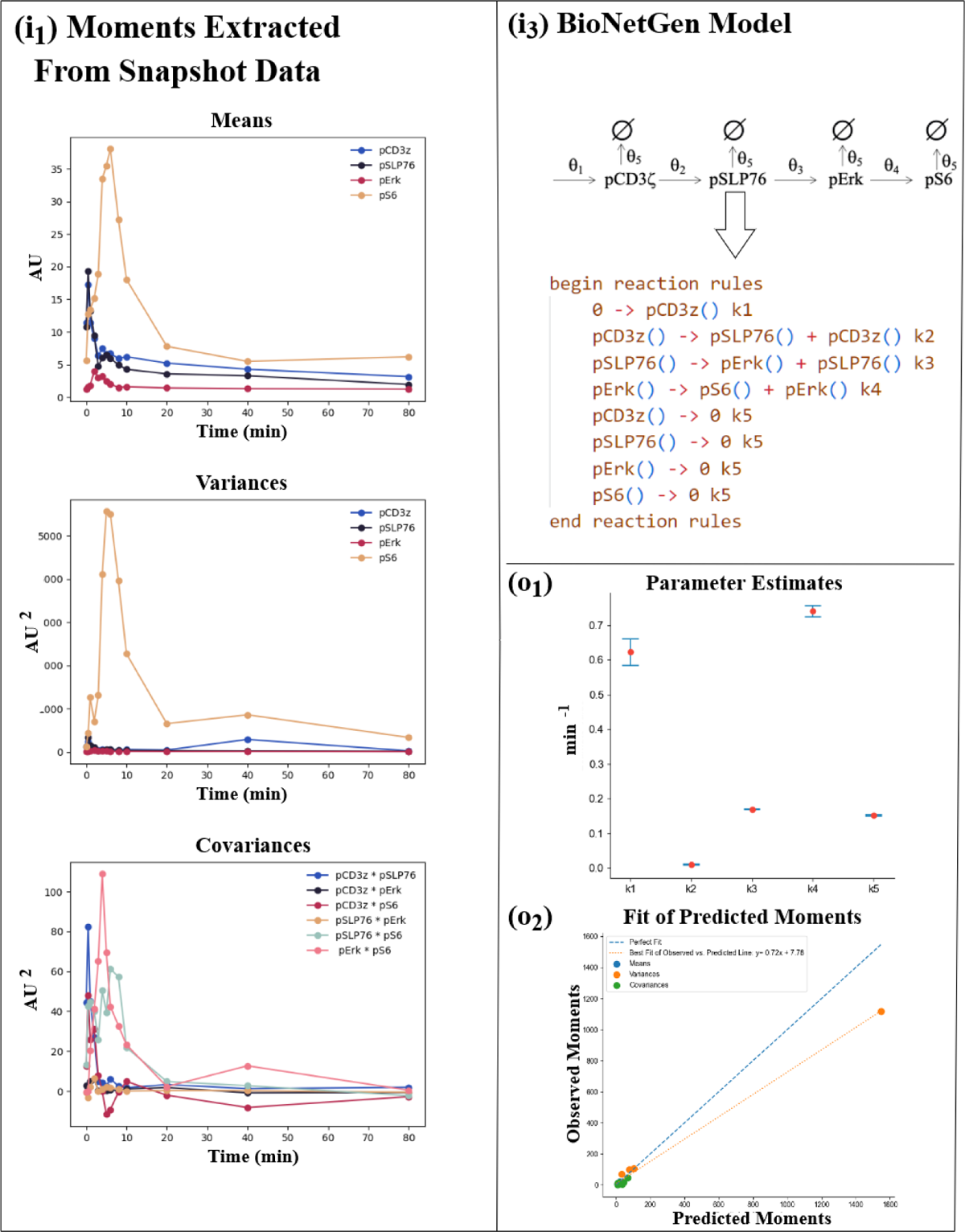
CD8+ T cell Example. The input and output results for fitting the CD8+ T cell CyTOF dataset are shown. The panel labels (i_1_), (i_3_), (o_1_), and (o_2_) correspond to those in Figure 1. Panel (i_1_) in Figure 3 shows the sample means, variances, and covariances of CD8+ T cell proteins calculated by BioNetGMMFit for the observed time points of snapshot data (supplied in the files i_1_ and i_2_). These are used for parameter estimation as part of the GMM procedure. (More information on GMM can be found in the Appendix.) Panel (i_3_) displays the reaction network and its corresponding reaction rules in the .bngl file. Each reaction rule represents a reaction within the model and has a parameter (rate constant) associated with it (e.g., k1). Using the observed moments (i_1_ panels) BioNetGMMFit uses PSO and GMM to estimate parameters of the model. (o_1_) and (o_2_) illustrate the results for a case where BioNetGMMFit performs 30 PSO estimates and produces a set of confidence intervals as well as a plot of the least GMM cost fit between the observed moments and predicted moments. The fit is performed with 4 protein means, 4 protein variances, and 6 protein covariances. Please note that the user is responsible for the consistency of units units in the input data since BioNetGMMFit uses the input value without ascribing about measurement units. For the CD8+ T cell data used above the input data were of the magnitude of fluorescence.

Once the model is defined, the hyperparameters configuration file supplies the PSO hyperparameters (i.e the number of particles, steps, and PSO coefficients) used for parameter estimation. For more information on the configuration file, a table of parameters is provided in the documentation on the GitHub page and in the Appendix. To estimate parameters within the BioNetGen model, PSO minimizes the GMM-weighted square euclidean distances of lower and higher order moments between the observed and model-simulated data [7]. Upon PSO convergence, estimates are provided and confidence intervals are computed. The plots of parameter estimates and moment fits provide additional insights into the fit of the mechanistic model. In Figure 3, for example, the confidence intervals around the parameter estimates reveal the ranges of possible values for each model parameter and their relative magnitudes. The estimation shows that the rates of production of pCD3*ζ* and of pS6 induced by pErk are the two dominant signaling reactions that occur at the early times (1-2 mins) when CD8+ T cells are stimulated. This result is consistent with related conclusions in [20] obtained using information theory. Nonetheless, models are imperfect, and their fits may be poor. To assess the model fit in such cases, we can refer to the “Fit of Predicted Moments” plot in Figure 3. This plot shows that the estimated model fails to accurately fit one of the protein’s variances, as indicated by a variance dot biased below the perfect fit line, indicating that further modifications in the model are needed to better fit the data. Such moment plots are essential for evaluating the parameter estimation and model fits that may be difficult to observe with only mean or bulk abundance differences.

BioNetGMMFit is currently available as a docker image for ease of portability across different operating systems, a compilable executable for those requiring high-performance computing capabilities (https://zenodo.org/record/7733865), and as a web demo (https://bngmm.nchigm.org/). Further details on how to use and compile BioNetGMMFit are available on the GitHub page (https://github.com/jhnwu3/BioNetGMMFit) and a comprehensive tutorial on using BioNetGMMFit can be found in the Appendix.

## 3 Results

We applied BioNetGMMFit to simulated datasets generated by models with known ground truth parameters to provide a reference point to evaluate its robustness, efficacy, and versatility. Specifically, we apply BioNetGMMFit to three simulated datasets, where each dataset is generated from a different model. Furthermore, we showcase the software’s functionality, which includes (1) facilitating estimation of model parameters from different combinations of moments for any rule-based model, (2) fixing subsets of model parameters both for simulation and estimation, and (3) forecasting moments at future time points. For each simulated dataset, we generated two sets of initial conditions from lognormal distributions as they have been observed across a variety of single-cell systems [24, 25]. To mimic the experimental constraint that “X” cannot be used to generate the observed snapshot data “Y” at time t, we generate another set of unobserved initial conditions that are used to obtain “Y” by evolving the BioNetGen models with ground truth parameters. This feature of simulating data with BioNetGen models when given a set of initial conditions is built into BioNetGMMFit. Figure 4 provides a diagram of this process.

**Fig. 4.**
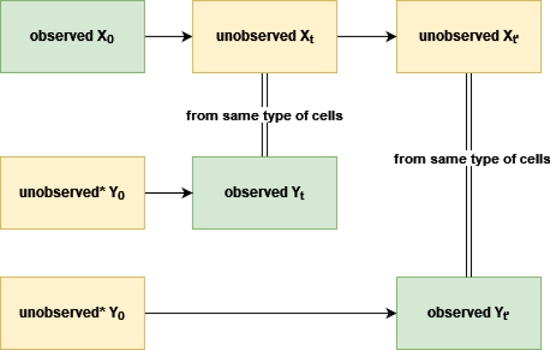
Data Generation Process: A fundamental constraint of time-stamped CyTOF data is that in order to measure a cell, it must be sacrificed. Therefore, in physical experiments, the initial conditions of cells observed at time *t* and *t^′^* are unknown. Similarly, cells observed at time *t*_0_, cannot be observed at future times *t* and *t^′^*. Here, we refer to such initial conditions and future states (shown in yellow) as “unavailable”. By contrast, parameter estimation only makes use of the “observed” data shown in green. Note that the “unavailable” data marked with an asterisk (*) must be generated in order to simulate data obtained from CyTOF experiments. In particular, one must simulate the initial conditions of all cells so that disjoint subsets of cells can be evolved to future times of interest. For ease of exposition, the subset index is not shown. We use the X and Y labels here in order to remain consistent with the inputs defined in Figure 1.

For more information on the simulated datasets themselves, please see the GitHub page (https://github.com/jhnwu3/BioNetGMMFit/tree/main/example) in their respective “X” and “Y” directories.

### 3.1 A Biochemical Reaction System with First Order Reactions

Here we investigate and show that BioNetGMMFit can successfully reproduce the ground truth model parameters for a system of first-order reactions model where six molecular species are arranged in a linear architecture and react via first-order biochemical reactions as illustrated by the reaction network in Figure 5. The reaction system can represent or can be generalized to represent a variety of sequential cell signaling processes such as phosphorylation of adaptor 14 3.1 A Biochemical Reaction System with First Order Reactions proteins such as DAP12 or Fc*ɛ*R1*γ* in NK cells [26] which are phosphorylated by Src family kinases or series of chemical modifications in signaling proteins in membrane-proximal signaling events in T cells[27]. We successfully estimate all 6 parameters of the model and showcase one of BioNetGMMFit’s important features: the ability to estimate parameters using different moments. We note that one advantage of GMM (see Appendix) is that it allows for the use of any number of moments in its objective function. BioNetGMMFit allows the choice of fitting three different combinations of moments of across a dataset: (1) first moments (means), (2) first (means) and second moments (variances), as well as (3) first (means), second (variances), and mixed moments (covariances) as shown in Figure 6.

**Fig. 5.**
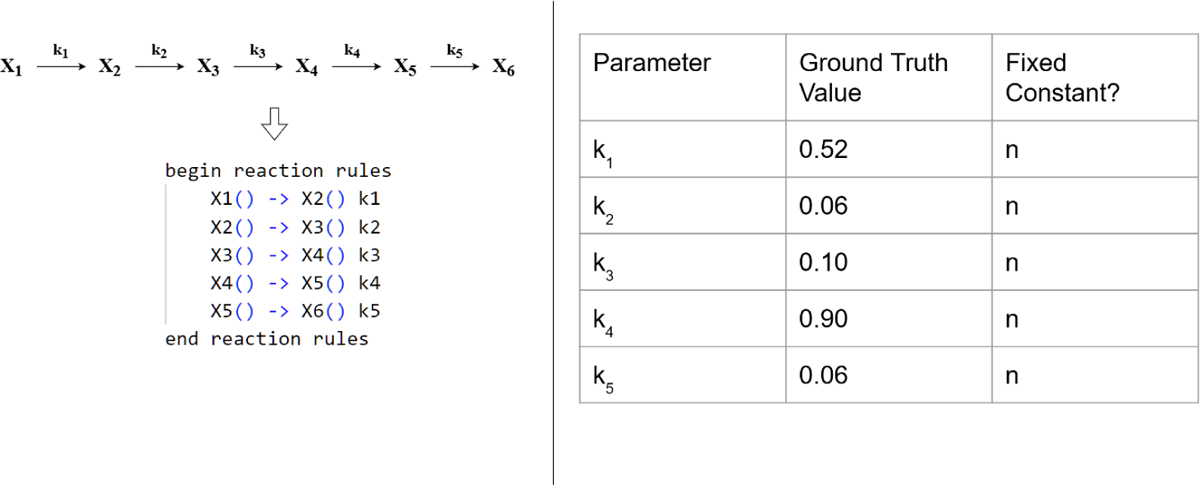
First-order reaction model: On the left, X_1_ through X_6_ represent protein species *^θ^*in a system of reactions where *−* represents a reaction in which species X*_i_* is changed into Click to show the PDF X*_j_* at the rate given by the corresponding *θ*. BioNetGMMFit is used to estimate the set of *θ*’s. Below the system of reactions, its BioNetGen reaction rules are displayed; these need to be defined and input to BioNetGMMFit to estimate the *θ*’s. Note that we use k’s instead of *θ*’s to define model parameters in BNGL. The table on the right contains the ground truth values.

**Fig. 6.**
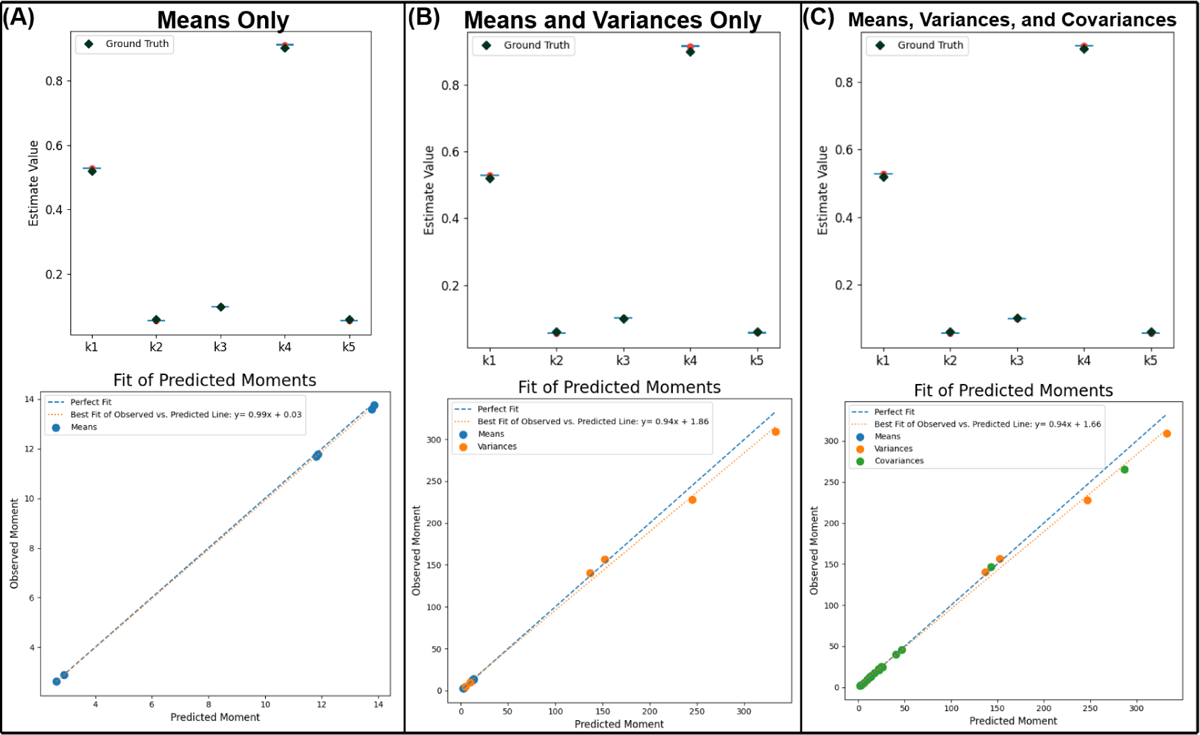
Parameter Estimates Using Different Combinations of Moments in the First-order reaction model: The left column (A) contains 95% confidence intervals of parameter estimates obtained using only the means of the snapshot data at time t=1.5 (top panel) and the comparison of the predicted means and the actual (observed) means (bottom panel). The middle column (B) contains analogous results by fitting the means and variances. Note the change in the scale of the values of the moments when the variance is included. The last column (C) on the right are the results obtained by fitting all the first, second, and mixed moments or more specifically all 6 means, 6 variances, and 15 covariances of the 6 proteins. This illustrates the flexibility available to the user to choose different subsets of moments for parameter estimation. In the analysis, the initial conditions and the simulated observed conditions at time t=1.5 both contained 10,000 cells each.

A key strength of this flexibility of moment choice in the cost function is the ability to adapt to different problem requirements. For instance, inclusion of higher order moments tends to improve parameter estimation in this first-order reaction model as shown in Figure 6 whereas the usage of variances and covariances in its fitting routine produces the tightest confidence intervals around its parameter estimates.

### 3.2 Model for Msn2 Induced Transcription in Yeast

Next, as mechanistic models often incorporate some form of nonlinearity, we show that we can successfully perform parameter estimation for a nonlinear gene regulatory model involved in stress response in yeast [28]. This is a simplified version of a commonly occurring gene regulatory motif in yeast. One of the key regulators of the stress response is the transcription factor Msn2 which resides in the cytoplasm in the resting state and is dephosphorylated in response to stress. Upon phosphorylation, Msn2 translocates to the nucleus, binds to stress-responsive elements (STRE), and induces gene expression of response proteins. The model is illustrated in Figure 7. This example explores a scenario where certain model parameters need to be fixed. Here we fixed two parameters, the death rate *p_d_* and the birth rate *p_b_* of the protein species *P*, reducing the total number of parameters to be estimated to four. The BioNetGen rules describing the model and the ground truth parameter values used to generate the synthetic data are shown in Figure 7. We analyzed its simulated observed snapshot data at time points 0, 1, and 5 minutes in its parameter estimation.

**Fig. 7.**
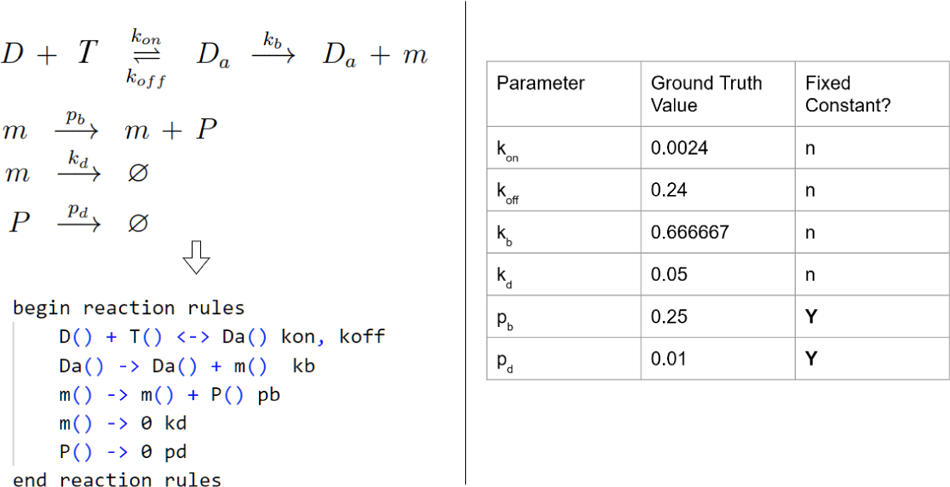
Msn2 Transcription Model: Model reaction network is shown on the left with its corresponding BioNetGen reaction rules defined below it. Ground truth model parameters and which ones have been fixed are shown in the table on the right. In this case, non-fixed model parameters labelled by “n” are estimated.

The parameter estimation results for the Msn2 transcription model are presented in Figure 8. Although the moment fits for both time points are not perfect, with an underfitted variance for one of the species, the confidence intervals around the estimated parameters fully encompass the ground truth parameters, indicating that BioNetGMMFit is capable of handling parameter estimation for nonlinear models.

**Fig. 8.**
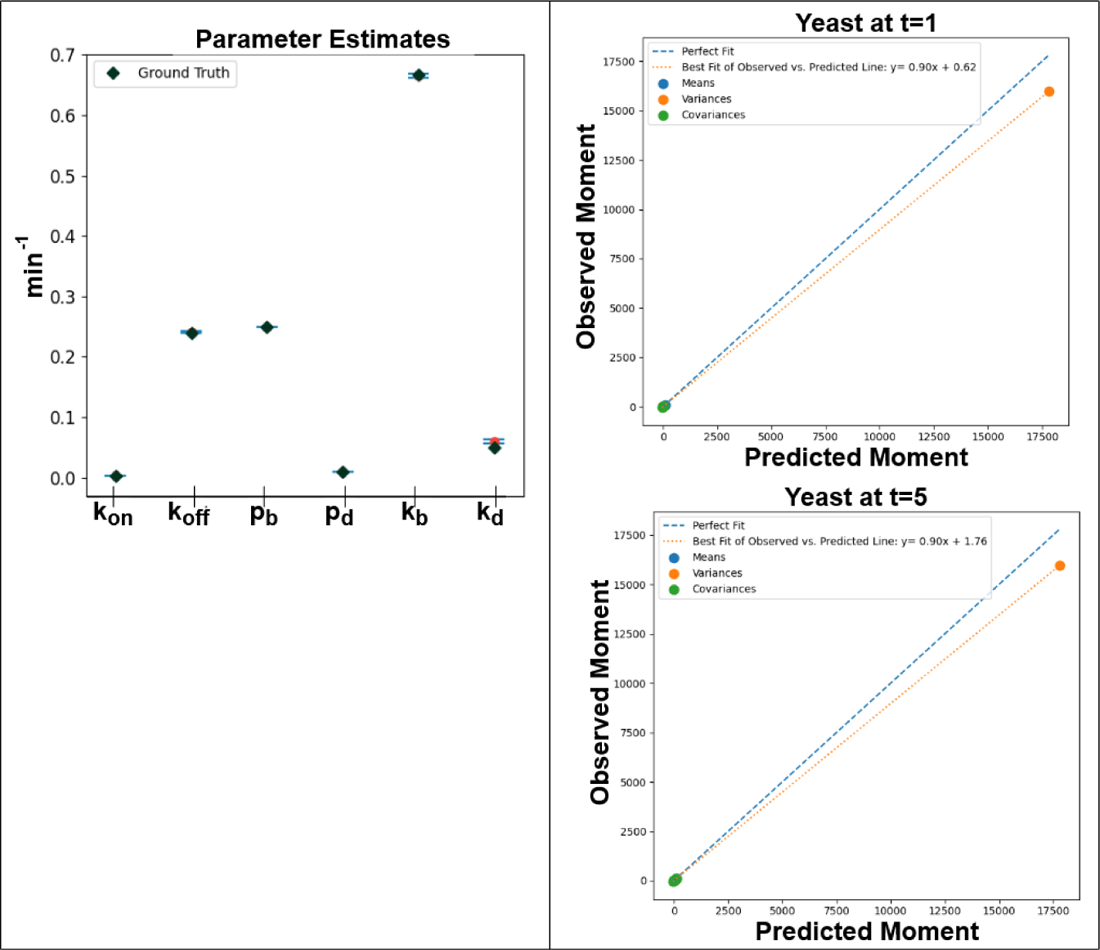
Msn2 Transcription Model Outputs: Model parameter estimate with confidence intervals on the left were generated by the moment fits for 2 different simulated observed time points, t=1 and t=5 minutes, on the right. In this case, 95% confidence intervals of parameter estimates shown on the left include the ground truth parameter values. Please note that the middle two x-tick marks, pb and pd are not estimated and are held constant. Using simulated data for the 5 species (i.e D, T_2_, m, etc.) in the ground truth model, 5 means, 5 variances, and 10 covariances across two time points were obtained and used for parameter estimation. All snapshot data used in the analysis contained 10,000 cells.

### 3.3 Vav1 activation kinetics in NK cells

As the final example, we consider a simplified model early time biochemical signaling kinetics (Figure 9) in mouse Natural Killer (NK) cells [29] to illustrate a case where identifiability issues prevent BioNetGMMFit from capturing the ground truth model parameters. The model describes the phosphorylation of a key signaling protein Vav1 by the kinase Syk bound to activating receptor-ligand complex and the dephosphorylation of phosphorylated Vav1 by the phosphatase SHP-1 bound to the inhibitory receptor-ligand complex. The nonlinear biochemical reactions in the model are akin to the zero-order ultrasensitivity model proposed by Goldbeter and Koshland [30]. We assume the abundances for the proteins Syk, Vav1, Syk-Vav1, SHP1, SHP1-pVav1, and pVav1 are measured at any time t. Following parameter estimation, we predict first, second, and mixed moments. The comparisons between the predictions and the simulated observed moments at different time points are shown in Figure 10. The three slopes shown in Figure 10, each near unity, indicate good agreement between the predicted moments (i.e. means, variances, and covariances) and the observed moments at each time point. Furthermore, using the model parameters estimated by BioNetGMMFit, we can *forecast* moments (i.e. predict moments) at times well outside of the given observational range (Figure 10). The nonlinear Vav1 activation model also presents a case where hyperparameter tuning is needed to generate more accurate parameter estimates. For instance, in Figure 10, increasing the number of particles and steps in the PSO routine improves the parameter estimates (i.e., brought them closer to the simulated ground truth). Despite improved estimates, this scenario illustrates a parameter estimation problem where other parameter estimates exist that minimize the objective function. Figure 10 shows that although the moment fits are excellent and the predictions closely match the observed and future unobserved data, the parameter estimates for k_2_ and k_5_ are unable to capture the ground truth values, even with a tuned hyperparameter configuration of 1,500 particles and 150 steps of PSO. In the subsection below, we provide further insights into the challenges of parameter estimation in rule-based models.

**Fig. 9.**
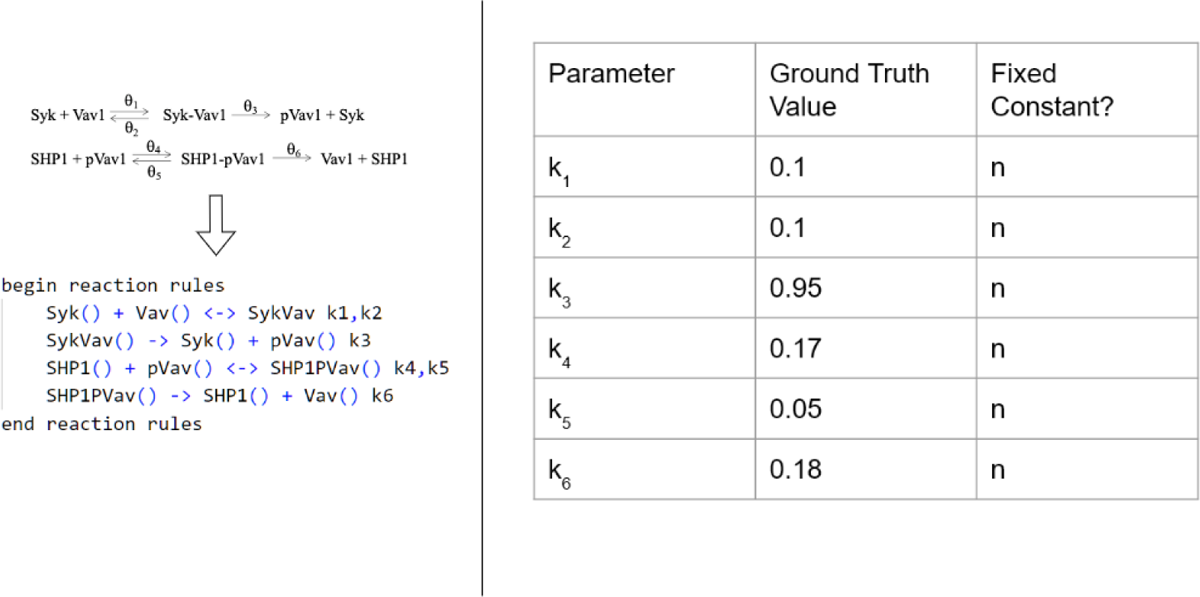
Vav1 activation model: The reaction network displayed on the left represents a simplified version of the phosphorylation-dephosphorylation kinetics of Vav1 in mouse NK cells. The corresponding reaction rules in BioNetGen are shown below. All the parameters, the set of rate constants *{θ_j_ }* or k*_j_* in the .bngl file and their ground truth values on the right are estimated using BioNetGMMFit.

**Fig. 10.**
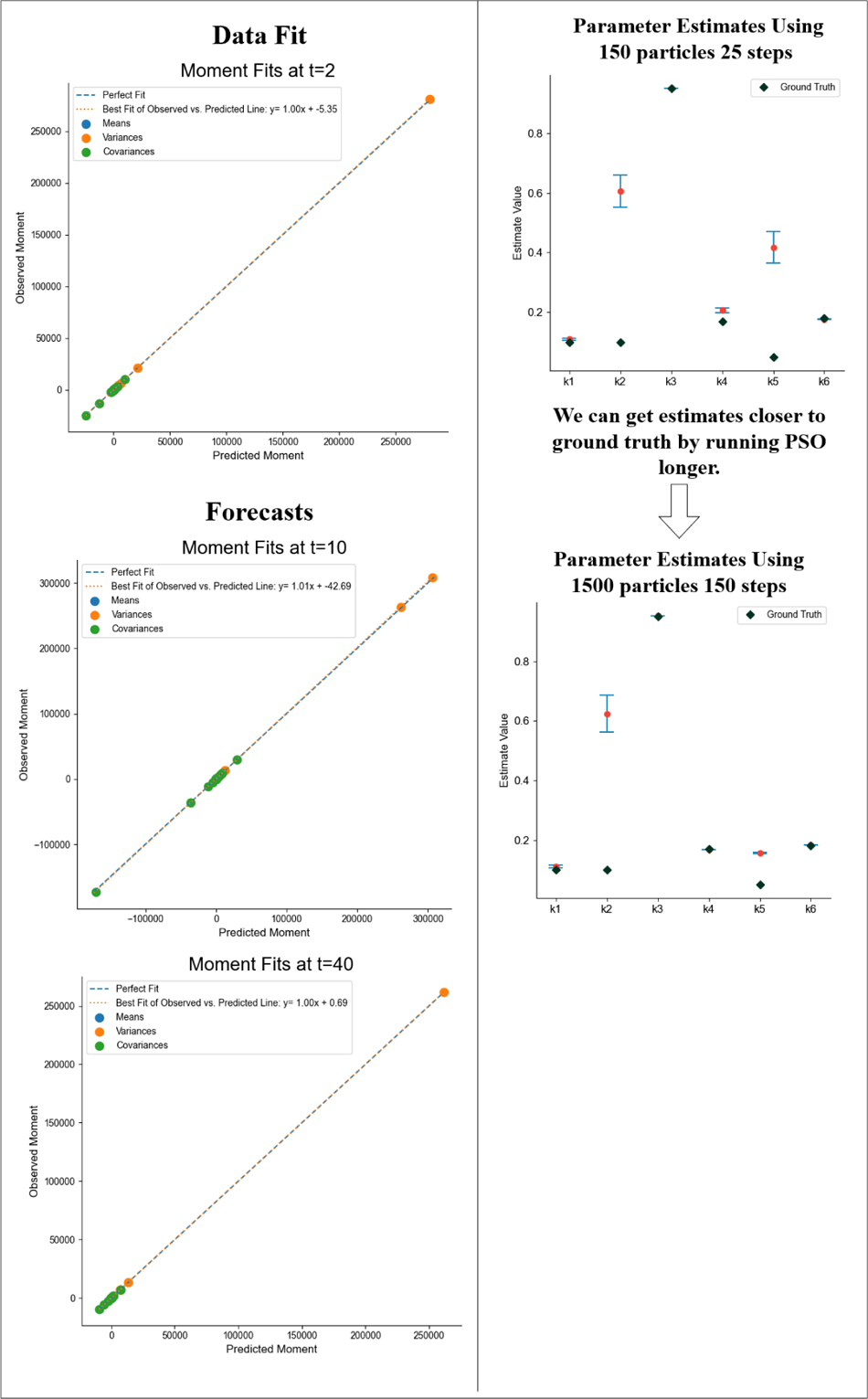
Results for the Vav1 Activation Model: The ground truth parameter values shown in Figure 9 are estimated by BioNetGMMFit using data from the ground truth model at *t* = 0.5 and *t* = 2. 5,000 cells at each time point were used in the analysis. The panels in the left column show the predicted moments calculated with estimated parameters plotted against the observed moments calculated using the ground truth model at various time points. The top figure shows the comparison at one of the input time points while the bottom two graphs are the “forecast” moments at t=10 and t=40. This shows that such parameter estimates can predict values at time points well into the future. The parameter estimates were derived from configuration 6 Protein A. Please see Table A2 in the appendix for more information on the PSO hyperparameter configurations used. On the right, we show improvements in estimates by tuning PSO hyperparameters and show parameter estimates for two different hyperparameter configurations. All the moment comparison plots in the left column were generated with parameter estimated obtained using PSO with 1500 particles and 150 steps. Note that the time units for this simulated dataset are arbitrary since BioNetGMMFit does not have units built into its plotting function. However, phosphorylation reactions as defined by the network in Figure 9 occur on the time-scale of seconds.

### 3.4 Practical Concerns in Estimation

As seen from the Vav1 activation model, cost functions for ODE models often possess multiple local minima and/or flat regions with small curvatures [7], that typically make estimation (i.e. the task of finding the best parameter values) challenging for any method. Although PSO mitigates these optimization problems to some extent, depending on the computational constraints, BioNet-GMMFit does not fully address the difficulties of such irregular cost functions, e.g., BioNetGMMFit can still yield estimates that reflect a local, rather than the global, minimum. One such example is shown in Figure 10 where only 150 particles and 25 steps were used for parameter estimation of the simulated Vav1 activation model. We intentionally chose a limited number of particles and steps to examine what could happen in a computationally constrained environment. As one might expect, the estimates for some paremeters were not close to their corresponding ground truth values; however (perhaps surprisingly), the estimates were able to accurately predict the observed first, second, and cross moments at later times as shown in Figure 10. However, when the number of particles and steps used are increased tenfold, parameter estimates are much closer to the ground truth, although prediction improves only slightly. Moreover, this dramatic improvement in estimation and a slight improvement in prediction demand higher computational costs as shown in Table A3 where there is a drastic increase in run time from configuration “6 Protein Time Points Set A” to “6 Protein Time Points Set A Extreme” in Table A3. As such, this raises an interesting question of “How much should one care about obtaining the best possible parameter estimates (as shown with *k*_4_ in Figure 10 for example), especially when computational resources may be limited and/or when the primary interest of the analysis may be prediction, not estimation?”

Furthermore, despite the potential for dramatic improvement in estimation, there is no guarantee that one will realize the desired improvement simply by using more particles and/or steps. We acknowledge the previous work of Sethna et al. [31], which used information theory to characterize parameter identifiability issues and model sloppiness; and the work of Sorger et al. [32], which used a Bayesian approach to compare competing models in the presence of partial identifiability. Although BioNetGMMFit is not designed to directly address those issues, our software does give users the ability to inspect a given model for identifiability concerns by constructing pairwise contour plots of the high-dimensional cost (aka objective) function. For each contour plot the x and y axes correspond to a pair of distinct rate constants. For example, pairwise contour plots can be generated for the nonlinear Vav1 activation model, where local minima were virtually indistinguishable from the ground truth, resulting in parameter estimates that were different from the ground truth (see Figure 10). By creating pairwise contour plots for k_2_ (or *θ*_2_) and k_5_, where PSO was unable to discern the ground truth values (see Figure 11), a striped pattern on the log-cost contour plot reveals a region of unidentifiability. In contrast, the well-estimated first-order reaction model has a convex log-cost shape (see Figure 11).

**Fig. 11.**
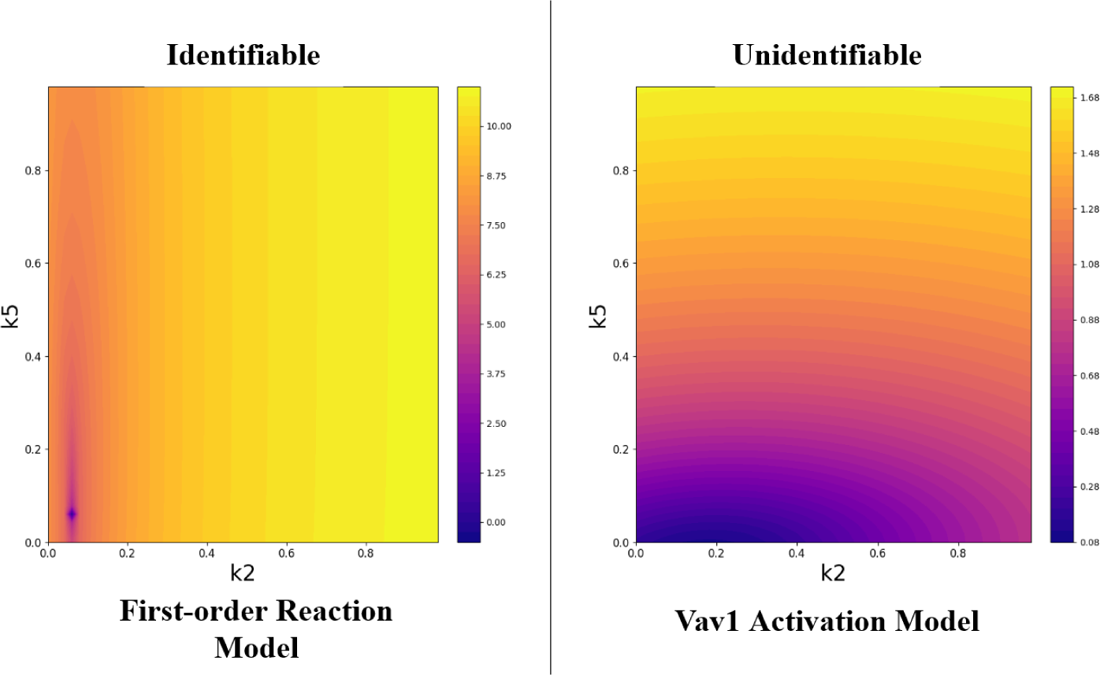
Contour Plots for Identifiability: Contour plots of the landscape of the cost function are shown as a function of k_2_ on the x-axis and k_5_ on the y-axis with all other model parameters held constant at the ground truth. The left contour plot is a pairwise contour plot of the fully “identifiable” first-order reaction system where a steeply convex cost landscape surrounds the ground truth parameters at the bottom left. The right pairwise contour plot is of the nonlinear Vav1 activation Model where a nonconvex cost landscape results in the parameter estimates shown in Figure 10. Note the big difference in the scale of the color values for the two plots. The colour bar is of the log-GMM cost that is derived by BioNetGMMFit with 6 means, 6 variances, and 15 covariances in its calculation.

### 3.5 Comparison with Other Software Tools

By combining the best aspects of CyGMM and BioNetGen, BioNetGMMFit provides several noteworthy features for the analysis of single-cell timestamped cytometry data that complements the currently-available parameter estimation software suite (i.e PyBioNetFit [18], Statistical Model Checking (SMC) [33], COPASI [22]). In particular, BioNetGMMFit offers the ability to efficiently analyze single-cell trajectories for a large number (*>* 1000) of cells, predict higher-order moments at future times, and simulate parameter estimation tasks with ground truth parameters. We detail the full capabilities and limitations of BioNetGMMFit below as well as how it compares to other pre-existing parameter estimation software packages. A brief summary of the software-related differences are shown in Figure 12.

**Fig. 12.**
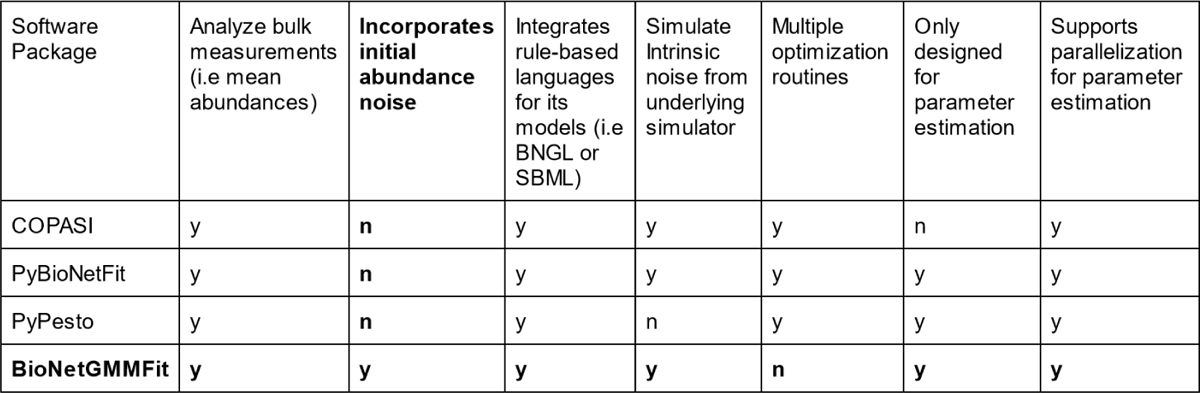
Comparison of BioNetGMMFit Features: Bulk measurements refer to measurements in which one obtains a single observed value for each species of reactants, at each time point. The observed value may be the mean or total abundance of the species at that time point. In contrast, single-cell measurements yield copy numbers or concentrations of each species for thousands of cells at each time point. This leads to what we refer to as initial abundance noise. The letter “y” indicates that the feature is available and “n” implies that the software is not designed to do it.

### Dealing Efficiently with Cell-Specific Initial Conditions

BioNetGMMFit accurately and efficiently estimates BioNetGen model parameters from cytometry snapshot data observed across many single-cell trajectories. It employs the same sophisticated minimization procedure as CyGMM [7] to estimate parameters from the analysis of higher-order moments. To the best of our knowledge, existing software are not designed to account for large variances in initial conditions. Thus, they have difficulty performing parameter estimation efficiently where (1) initial abundance noise is known to play a role and (2) the data consists of large numbers of different single cells (where each cell has its own set of trajectories). For example, when the estimation depends on the analysis of data from a large number of single cells (as opposed to bulk measurements averaged over cells), existing software is eithernot designed to handle such data [18, 33], or is less user-friendly, or is computationally inefficient [14]. In many cases, when attempting to perform parameter estimation across a combination of moment statistics, inefficiencies related to file input and output occur. For instance, in using BioNetGen’s simulator, each cell has its own BioNetGen file containing a bulk measurement such as average protein abundances. Therefore, to perform parameter estimation across a combination of different measurements (i.e means, variances, and covariances) with BioNetGen only [14], the user must repeatedly read from multiple data files in its parameter estimation. In PyBioNetFit [18], only a single set of cell conditions per time point are used for parameter estimation. Our method allows users to input easily several time-stamped snapshot files containing abundance data from thousands of cells and store them directly in memory for quick simulations. See Table A1 below for each dataset’s cell count used for parameter estimation. While the comparison is not exact as the models and objective function (i.e BioNetGMMFit’s incorporation of higher-order moments) analyzed were different, we show that we can attain run-time performance comparable to that of PyBioNetFit [18]. In the analysis of the Vav1 activation model in Figure 9 with 5,000 single cells, the run time was approximately 1.9 hours on a 16-core processor for 30 repeated parameter estimates, which is in the range of run time using the SBML simulator in PyBioNetFit for a similar number of cores [18]. To provide some context on run time, we compare with a method such as SMC [33]: while again the comparisons are not exact as the methods and models are different, our run times are comparable to their 4.2 hours for parameter estimation. Due to C++ parallelization with OpenMP, our software is capable of linearly scaling with increasing cell count, thereby reducing run times for large numbers of cells. Table A3 provides run-time details concerning the number of cells and model parameters.

### Estimating with Higher-Order Moments

Due to their design intended for bulk measurements, existing software such as PyBioNetFit [18], COPASI [22], and PyPesto (from the PETab suite) [34] for rule-based parameter estimation are not specifically designed to fit higher moments. While possible to do a plethora of file input and output to generate multiple parameter distributions per set of initial conditions and generate moment statistics using PyBioNetFit or BioNetGen, for a large number of cells, parameter estimation becomes very user unfriendly and computationally taxing. In contrast, BioNetGMMFit directly supports three levels of analysis corresponding to using an increasing number of moments. First, one can choose to estimate parameters using only the means (i.e. first moments). Second, one can use both means and variances or third, use means, variances, and covariances. Results for moments with different levels of fitting are shown with its different columns in Figure 6. Other more mature softwares such as COPASI [22] offer a wide variety of other features such as an executable graphical user interface, the ability to export ODEs from these rule-based languages, as well as the support of a wide range of optimization routines. For optimization routines involving smaller and simpler (i.e linear instead of nonlinear) models where large initial abundance noise is not of concern, it is preferable to use COPASI [22], PyBioNetFit [18], and PyPesto [34] due to the availability of less computationally expensive gradient-based optimization routines. However, in such simple parameter estimation tasks, BioNetGMMFit performs functionally the same due to its flexibility of cost functions where the user can simply choose to use only the observed first moments.

## 4 Discussion

We present a new software: BioNetGMMFit, that greatly enhances a researcher’s ability to explore a wide range of mechanistic models while using higher-order moments to leverage the information in single-cell data. In particular, BioNetGMMFit provides accurate estimation of model parameters, and reports valid confidence intervals for each estimated parameter as well. Furthermore, BioNetGMMFit can easily handle data observed across multiple time points, and it can predict protein abundances at future times.

A key feature of BioNetGMMFit, which benefits from its integration with libRoadRunner’s simulators, is its ability to easily simulate protein abundances at scale (i.e. across multiple time points, across many single cells, and under a wide variety of different biochemical signalling models). Moreover, because such simulations can now easily be carried out at scale, the ability to tune hyperparameters has become a much more accessible task.

Hyperparameter tuning involves selecting the optimal set of hyperparameters of the optimization routine to achieve the best parameter estimates that minimize the cost function and enable optimal prediction from one’s model. For example, in the case of PSO, there are five hyperparameters to tune, including the number of particles, the number of steps, and three coefficients that affect how the particle swarm moves in the space of model parameters. To address this challenge, we use a simulated reference point of the “optimal” or ground truth parameters for a BioNetGen model, which can provide insights into the performance of an optimization routine’s hyperparameter configuration. A practical way of determining a suitable set of hyperparameters when analyzing experimental data is as follows. If one wishes to fit a model to the data, one can simulate a ground truth version of the model separately and adjust the hyperparameters so that BioNetGMMFit estimates the ground truth values to desired accuracy. Then, one can use these newfound hyperparameters to fit the model to the experimental data.

### Limitations of BioNetGMMFit

BioNetGMMFit is currently limited to parameter estimation for well-mixed mechanistic models describing deterministic and stochastic kinetics. Unlike PyBioNetFit [18], other forms of simulation such as NFsim [35] and spatial modeling [36, 37] are not supported. Furthermore, BioNetGMMFit does not support any optimization algorithms other than PSO. Additionally, the lack of Python support may make it difficult to integrate it with pre-existing workflows that rely heavily on Python. While there exists a C++ static binary that has been tested on Ubuntu Linux, compilation with libRoadRunner C++ libraries can be a nontrivial process on other Linux distributions. Thus, BioNetGMMFit is most readily accessible through Docker, which although accessible across all major operating systems, still requires the use of third-party virtualization software. These software weaknesses will need to be addressed in future iterations of the software.

## Data Availability

The dataset(s) supporting the conclusions of this article is(are) available in the BioNetGMMFit/example repository, [https://github.com/jhnwu3/BioNetGMMFit/tree/main/example].

- Project name: BioNetGMMFit
- Project home page:https://github.com/jhnwu3/BioNetGMMFit
- Archived version: N/A
- Operating system(s): Platform independent through Docker
- Programming language: C/C++
- Other requirements: Docker, if Compiling see GitHub
- License: MIT License
- Any restrictions to use by non-academics: N/A
- Other Locations of Access: https://hub.docker.com/repository/docker/jhnwu3/bngmm/, https://zenodo.org/record/7733865, https://bngmm.nchigm.org/

Experimental CD8+ T cell data can be found here (https://dpeerlab.github.io/dpeerlab-website/dremi-data.html).

## Acknowledgments

Special thanks to Dr. Jim Faeder, Dr. Ali Sinan, and Dr. Lucian Smith for their advice in the development of this software. This work is supported by the NIH awards R01-AI 143740 and R01-AI 146581 to JD, and by the Research Institute at the Nationwide Children’s Hospital.

## Author Contributions

Every author contributed to the writing and approval of this manuscript. John Wu and William CL Stewart contributed to the software development and design of BioNetGMMFit.

## Appendix A

### A.1 GMM Primer

Here, we give a short and simple exposition of MOM (method of moments) to motivate and describe its extension: GMM (generalized method of moments). Consider data *D*_1_*, …, D_n_*, where each *D_i_* is independent and identically distributed *N* (*µ, σ*^2^). MOM equates sample moments such as *D* and *D*^2^ to their corresponding expected values (*µ* and *σ*^2^ + *µ*^2^). Under a mild set of conditions, these equations can be solved yielding consistent estimates (*µ*^ and *σ*^) of the unknown parameters (*µ* and *σ*). Now, to illustrate how important MOM can be, let’s consider another simple example. Suppose we have data *Z_i_* where each *Z_i_* is distributed *Beta*(*α, β*). We can define the *j*th sample moment (denote *m_j_*) as 1*/n* ^L^ *Z_i_^j^*. Now, using the first and second sample moments, MOM yields the following equations: *Z* = *α/*(*α*+*β*), and *Z*^2^ = *α/*(*α*+*β*)^2^[*β/*(*α*+*β* +1)+*α*].

With two equations, and two unknowns, estimates *α*^ and *β*^^^ are the closed form solutions for *α* and *β* given in terms of the sample moments *m*_1_ and *m*_2_. Note that for this simple example one must resort to numerical methods to find the maximum likelihood estimates of *α* and *β*. Furthermore, as there are an infinite number of moments (and sample moments), the decision to use *m*_1_ and *m*_2_ is somewhat arbitrary. In practice since expectation values of lower order moments have lower errors for finite size datasets, lower order moments (e.g., means, variances, covariances) are usually considered for parameter estimations.

GMM (generalized method of moments) is a generalization of MOM that yields consistent estimation of *θ* (i.e. a vector of unknown parameters) when the number of moment equations (*aka* moment conditions) is larger than the dimension of *θ* (i.e. the number of unknown parameters). In this manuscript we have snapshot data observed at multiple time points, but for simplicity, let’s consider snapshot data X observed at time 0 and snapshot data Y observed at time *t*. The sample moments in Y are then compared to their corresponding expected values, which are approximated by first evolving X to time *t* for a given *θ* [denoted h(X;*θ*, t)] and then computing the sample moments of h(X;*θ*,t)]. Because the system of equations is often overdetermined, there typically isn’t a solution (i.e. no value of *θ* makes the difference between sample moments exactly zero). Therefore, the GMM approach advocates finding the value of *θ* that *minimizes* the distance between the sample moments of Y and the sample moments of h(X;*θ*,t)]. Let’s denote this *k*-dimensional vector of differences by Δ*m*(*θ*), so that the distance between sample moments is simply [Δ*m*(*θ*))*W* (Δ*m*(*θ*)]*^T^*. For example, when *W* is the [*k* x *k*] identity matrix, the GMM cost is the usual geodesic distance in *R^k^* which also corresponds to a *least squares approach* and has been shown to perform poorly compared to the GMM cost that uses the optimal *W* [7]. In practice, one often begins with *W*_1_ equal to the identity matrix, and then minimizes [Δ*m*(*θ*)]*W*_1_[Δ*m*(*θ*)*^T^*] to obtain the first estimate of the unknown parameters, denoted *θ*^^^_1_. Then, dance with (Hansen et al. [4]), the next iterate *θ*^^^_2_ is obtained by minimizing [Δ*m*(*θ*)]*W*_2_[Δ*m*(*θ*)*^T^*]. This process can be repeated for any number of steps; but for the standard two-step estimator, one simply stops at *θ*^^^_2_. Note that *W*_2_ involves matrix inversion, which is often done numerically. Finally, in order to find the value of *θ* that minimizes [Δ*m*(*θ*)]*W* [Δ*m*(*θ*)*^T^*], one should perform an efficient search over the parameter space. In particular, for the problems discussed in this manuscript (and for most problems in general), a brute force grid search is computationally infeasible. Fortunately, there are many such optimization routines to choose from, and we find that particle swarm optimization (PSO), which is amenable to parallel computation, works well even when the dimension of *θ* is large, and the distance function [Δ*m*(*θ*)]*W* [Δ*m*(*θ*)*^T^*] is multi-modal.

For a practical tutorial on GMM, please see our Python tutorial (https://github.com/jhnwu3/BioNetGMMFit/blob/main/example/gmmtutorial.ipynb).

### A.2 BioNetGMMFit Tutorial for CD8+ T cell Dataset

The data we are looking at is a set of snapshot data files across different time points. Such snapshots across time are shown below as box plots in Figure A1.

**Fig. A1.**
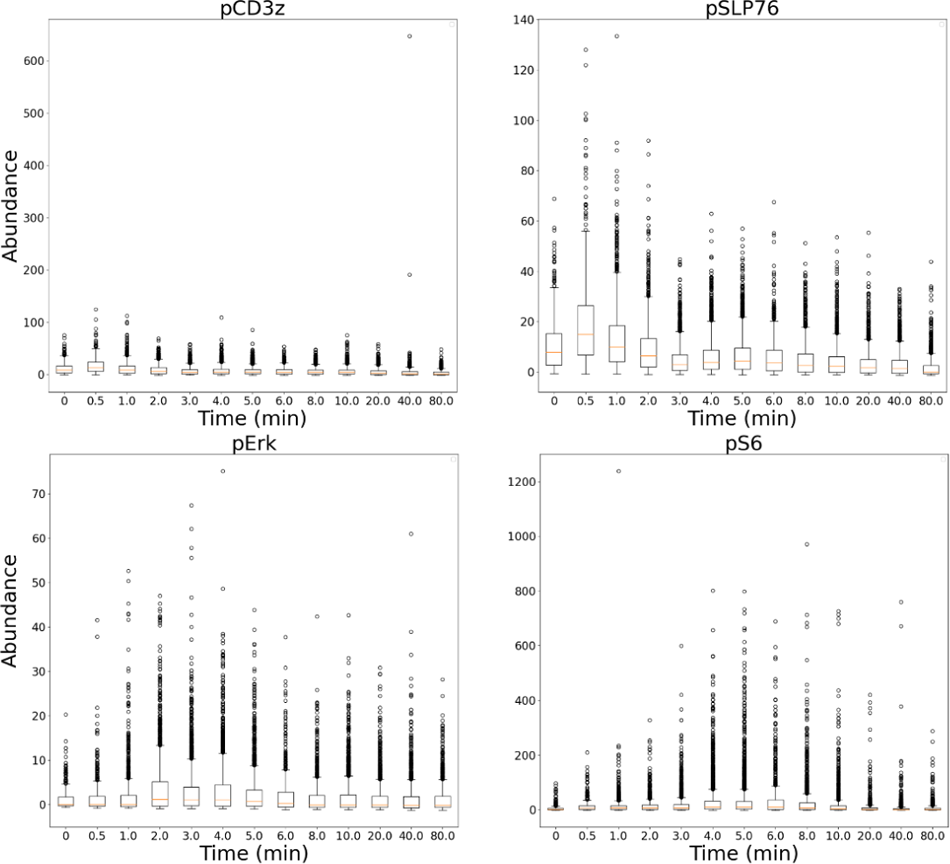
CD8+ T cell Snapshot Data: Black circles indicate points outside of the quartiles i.e outliers in the data. Please note that the x-axis is not relative to scale of time points.

First, let us consider the command line call in Figure A2

**Fig. A2.**
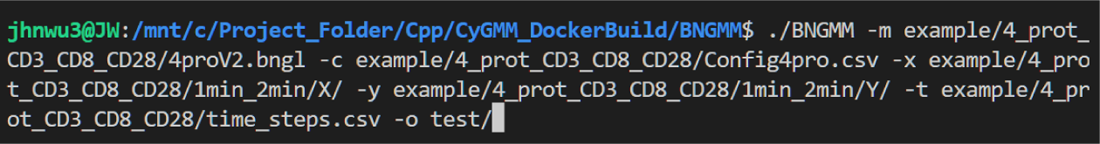
BNGMM Command Line Call

Inside this call, there are multiple required command line arguments that are explained in Figure A3 below.

**Fig. A3.**
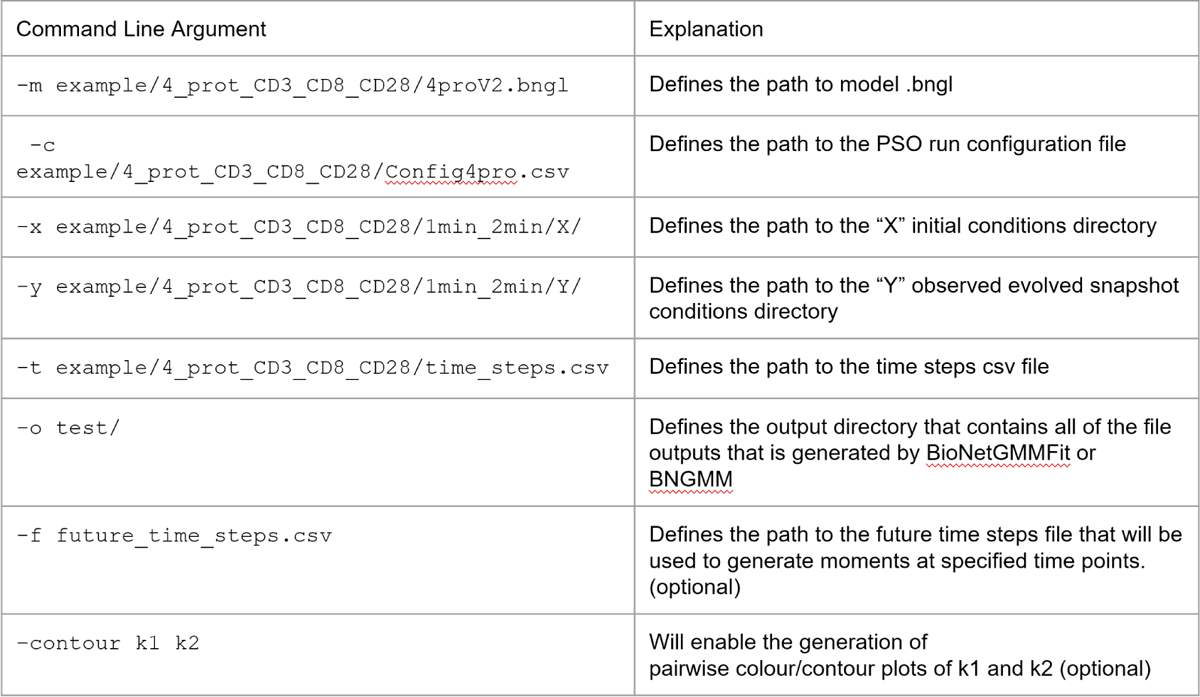
BNGMM Command Line Arguments

Taking a look at the .bngl file in Figure A4, there are several important caveats. The observables list also define the order in which data must be organized in its respective data csv files where the top observable matches the first column of the csv data files. Likewise, the order of the list of the parameters matches the vector that will be estimated. Please also note that the .bngl file has to write to SBML for BioNetGMMFit to work.

**Fig. A4.**
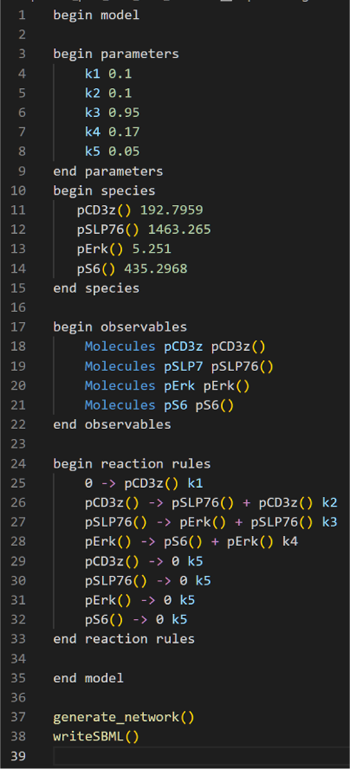
CD8+ T cell BioNetGen Model File

Zooming in on the data directories and time steps file, please note that the X directory contains specifically a csv file containing the initial conditions of the snapshot of CD8+ T cells at time t=1 minute. Again, the Y directory contains other csv files of the observed snapshot data at future time points. In the case of usage, the user should make sure that each Y directory file is named the same and has the time point number at the end of the file name as BioNetGMMFit reads in each file alphanumerically such that file with the smallest time point in its name corresponds to the first future time observed in the time steps csv file. A diagram of X and Y directories along with a set of time steps from the time steps file is shown in Figure A5. In this scenario, since there are only two time points, the initial conditions is considered to be at time t=1.0 minutes and the final observed conditions is at time t=2.0 minutes, and this corresponds to only one file being in the initial conditions directory and the other file being in the other observed conditions directory.

**Fig. A5.**
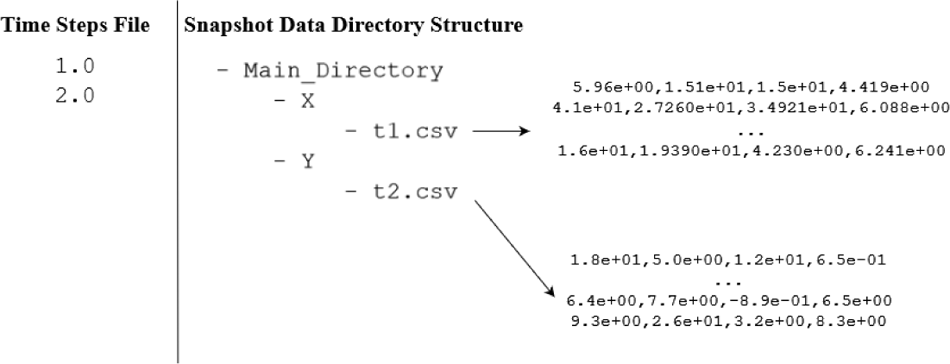
BioNetGMMFit X and Y directory structures

A BioNetGMMFit parameter estimation run is then characterized by its configuration file, which defines several key PSO hyperparameters such as the number of particles and steps. An example is given in Figure A6. All parameters in the configuration csv file are explained in Figure A7.

**Fig. A6.**
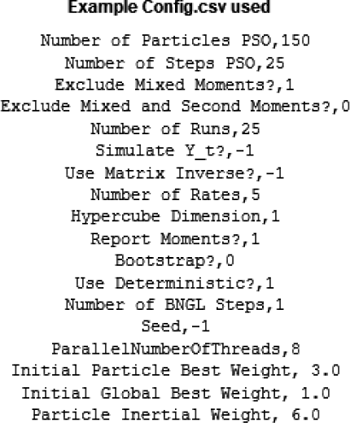
BioNetGMMFit Configuration File Example

Once the parameter estimation run begins, the command line program will output information such as the weight matrices computed from the snapshot data and observed moments. The number of weight matrices correspond to the number of snapshot csv files in the Y directory. Once the parameter estimation finishes, estimates and their corresponding costs are outputted as shown in Figure A8. Additionally, plots of moment fits and their 95 % confidence intervals are also generated as displayed in Figure A9. Should the user want to, forecasting can also be performed with the mean parameter estimate of their model using “-f futureTimesFile.csv” as part of the command line call.

**Fig. A7.**
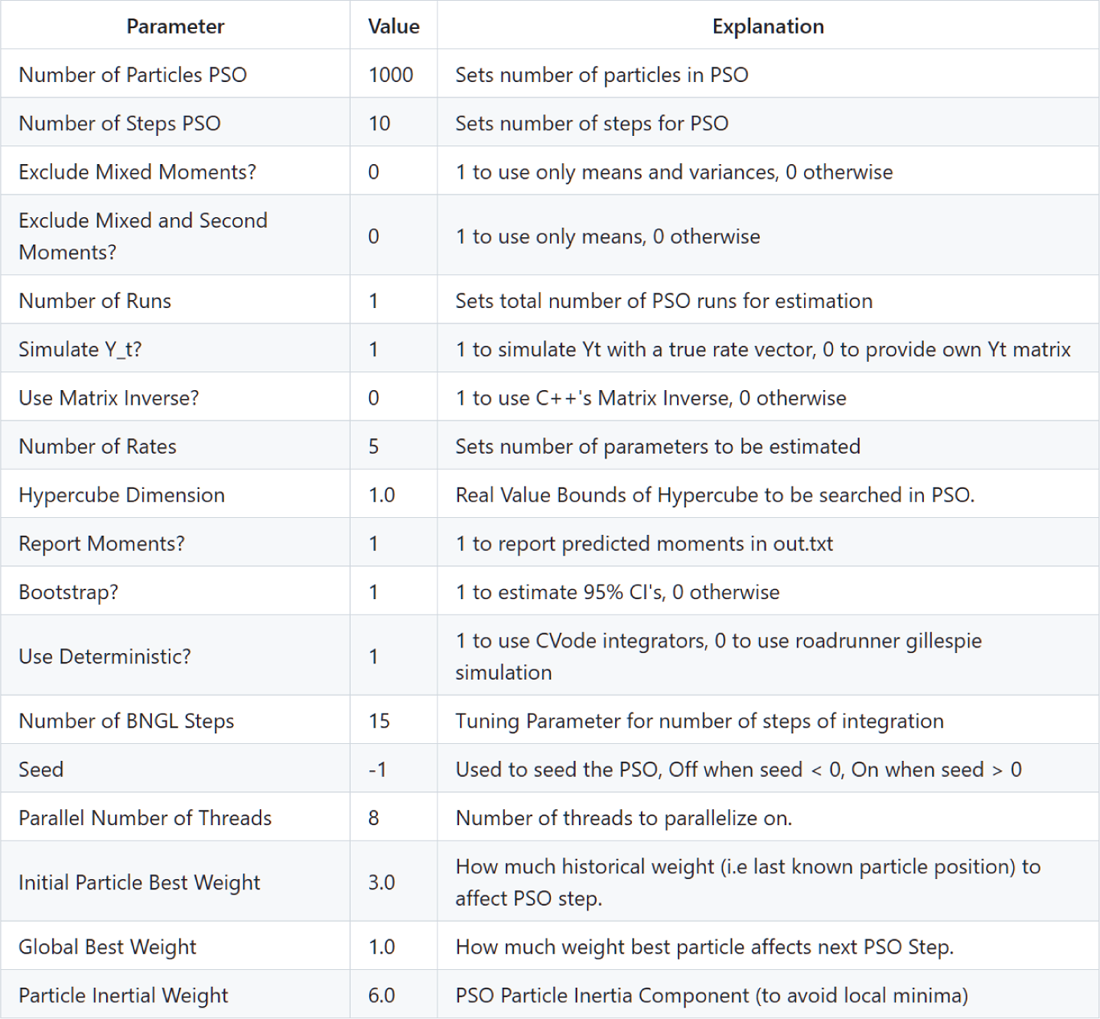
BioNetGMMFit Configuration File Parameters Explained

**Fig. A8.**
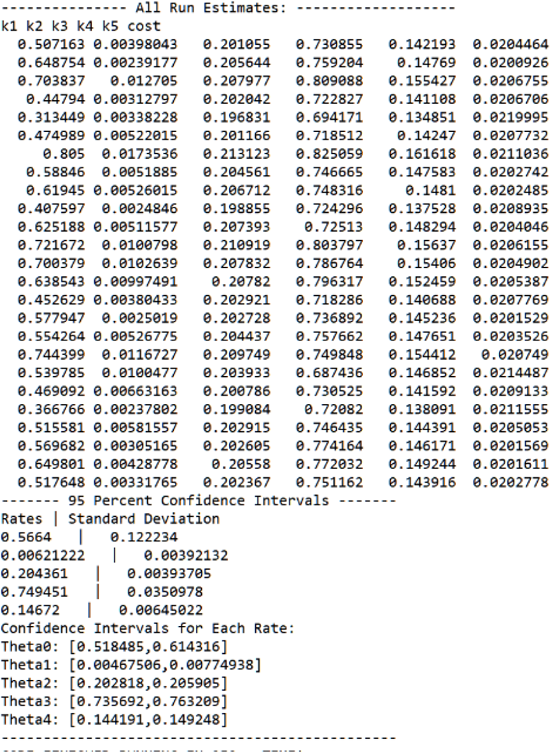
BioNetGMMFit Parameter Estimates: Generated by the C++ executable, the “All Run Estimates” contains the labels of the model parameters with each row being a parameter estimate and its associated GMM cost.

**Fig. A9.**
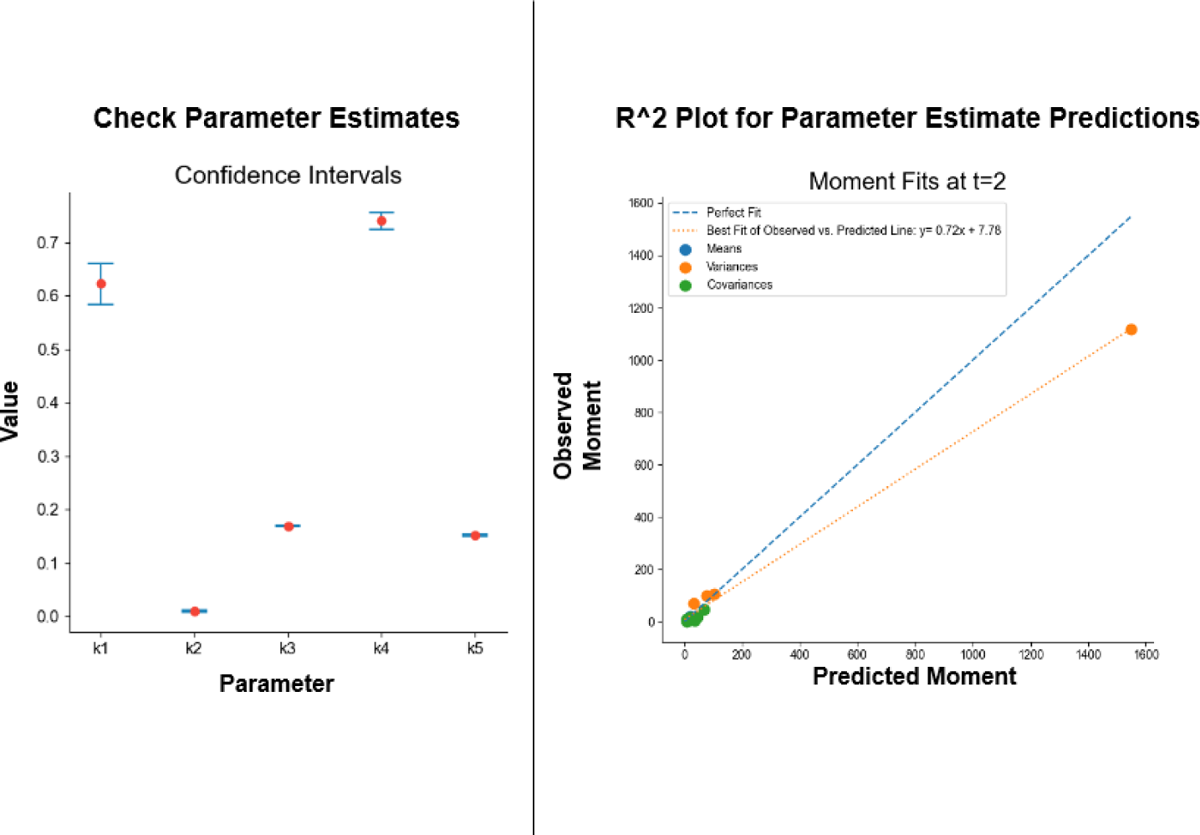
BioNetGMMFit Graphical Outputs

Again, for more information and details on the example along with a UI parameter estimation counterpart, please see the GitHub repo (https://github.com/jhnwu3/BioNetGMMFit).

### A.3 Computational Costs

Run times may vary drastically depending on the time points, models, and numbers of single cells being analyzed. To provide some idea of computational costs, we profiled the CD8+ T cell signaling model and the nonlinear signaling model shown in Figure 3. The number of model parameters, PSO hyperparameters, and dataset characteristics are shown in Tables A2 and A1 while their respective run times are shown in Table A3. Please note that each run is considered a PSO estimate. To generate confidence intervals, more than one run is required while generally 30 or more runs are preferred. All run times are rounded to the nearest second and all runs were performed using a 16-Core AMD EPYC 7302 processor with 32 threads and a maximum clock speed of 3.3 GHz. Only 30 threads were used for testing.

**Table A1.** Datasets Used For Estimation: The first row contain the labels of the model as well as its respective PSO hyperparameters and dataset contents used in BioNetGMMFit. Only two datasets are benchmarked, the 4 Protein CD8+ T cell and the simulated nonlinear Vav1 activation dataset. Each letter “A”, “B”, and “C” in the “6 Protein Time Points Set” columns indicate one unique set of time points of snapshot data used in the parameter estimation for the simulated 6 protein model. The “Extreme” term in the last column represents a ten-fold multiplicative increase of particles and steps in PSO. For instance “6 Protein Time Points Set A Extreme” is the PSO hyperparameter configuration that improves the parameter estimates from “6 Protein Time Points Set A” as shown in Figure 10. Table A2 provides more information regarding the PSO hyperparameters used for parameter estimation. The following row is the number of single cells in the initial conditions (i.e first time point) that are used for simulation by the BioNetGen model for estimation. The next row is the number of cells observed in the following time points. Cell counts remained constant across all time points in the simulated 6 protein Vav1 activation dataset.

| Hyperparameter | CD8+ T cell Model | 6 Protein Time Points Set A | 6 Protein Time Points Set B | 6 Protein Time Points Set C | 6 Protein Time Set A Extreme |
| --- | --- | --- | --- | --- | --- |
| # of single cells Used for Estimation | 719 | 5,000 | 5,000 | 5,000 | 5000 |
| # of single cells Observed | 653 | 5,000 | 5,000 | 5,000 | 5000 |
| Time Steps | 1.0, 2.0 | 0.0, 0.5, 2.0 | 0.0, 0.5, 2.0, 7.0 | 0.0, 0.5, 2.0, 10.0, 20.0, 30.0 | 0.0, 0.5, 2.0 |

**Table A2.** PSO and Model Parameters: Again, the first row indicates the model as well as its respective PSO hyperparameters and dataset contents. In this case, only two BioNetGen models were used, the 4 Protein CD8+ T cell model and the simulated 6 protein Vav1 activation model. The “Extreme” term in the last column represents a ten-fold multiplicative increase of particles and steps in PSO. For instance “6 Protein Time Points Set A Extreme” is the PSO hyperparameter configuration that improves the parameter estimates from “6 Protein Time Points Set A” as shown in Figure 10.

| Hyperparameter | CD8+ T cell Model | 6 Protein Time Points Set A | 6 Protein Time Points Set B | 6 Protein Time Points Set C | 6 Protein Time Set A Extreme |
| --- | --- | --- | --- | --- | --- |
| Model Parameters | 5 | 6 | 6 | 6 | 6 |
| Particles | 150 | 150 | 150 | 150 | 1500 |
| Steps | 25 | 25 | 25 | 25 | 150 |

**Table A3.**
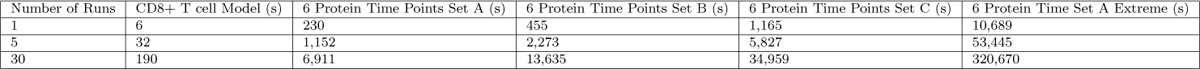
Run Times for Different Model Configurations: The 6 Protein Time Points Set A, B, and C columns show the change in run time with respect to the change in time steps used in Table A1. 6 Protein Extreme provides some insight into how long the run time of an extreme number of particles and steps used in PSO. One run is considered to be one PSO estimate. More than one PSO estimate is required for confidence intervals.

Although their asymptotic relationship may differ depending on the BioNetGen model, increasing the particle swarm’s number of particles and step count generally has a linear effect on run time where a 10*×* in particles tends to lead to a 10*×* in run time. In terms of data inputs, increasing the number of cells has a linear relationship with run time. More complex models have a dramatic effect on simulation time where increasing the number of parameters and species in a system increases run time in a nonlinear fashion. Similarly, as ode-solvers must approximate solutions, increasing the time evolutions tend to have a nonlinear relationship with run time. Fortunately, as BioNetGMMFit is parallelized through OpenMP, users can reduce run time by increasing the number of cores accessed by BioNetGMMFit.

